# Long noncoding RNAs and mRNAs profiling in ovary during laying and broodiness in Taihe Black-Bone Silky Fowls (*Gallus gallus domesticus Brisson*)

**DOI:** 10.1101/2023.08.17.553793

**Authors:** Yuting Tan, Yunyan Huang, Chunhui Xu, Xuan Huang, Zhaozheng Yin

## Abstract

Broodiness is one of the important factors affecting poultry egg production, with the black-bone Silky (*Gallus gallus domesticus Brisson*) exhibiting strong broodiness behavior. Changes in ovarian signaling can offer insights into the mechanisms influencing broodiness. However, research comparing the characteristics of long non- coding RNAs (lncRNAs) in the ovaries during the broody chickens (BC) and high egg- laying chickens (GC) groups of poultry is still limited. In this study, we employed RNA-seq to analyze the ovarian transcriptomes (lncRNAs and mRNAs) of eight brooding and high egg-laying Taihe Black-Bone Silky Fowls (TBsf). We identified a total of 16,444 mRNAs and 18,756 lncRNAs, among which 349 mRNAs and 651 lncRNAs (P < 0.05) showed significantly different expression (DE) between the BC and GC groups. Additionally, we identified the cis-regulated and trans-regulated target genes of differentially abundant lncRNA transcripts and constructed an lncRNA- mRNA trans-regulated interaction network associated with ovarian follicle development. Gene Ontology (GO) and Kyoto Encyclopedia of Genes and Genomes (KEGG) annotation analyses revealed that DE mRNAs and target genes of DE lncRNAs were associated with neuroactive ligand-receptor interaction, CCR6 chemokine receptor binding, G-protein coupled receptor binding, Cytokine-cytokine receptor interaction, and ECM-receptor interaction pathways. Our study provides a catalogue of lncRNAs and mRNAs related to ovarian development and constructs a predicted interaction network of differentially abundant lncRNAs and Differentially Expressed Genes (DEGs) in TBsf, thus enhancing our understanding of the interactions between lncRNAs and genes regulating brooding behavior.

**Author summary:** In this study, we performed transcriptome sequencing analysis of the ovaries from BC and GC groups in TBsf. A total of 349 significantly differentially expressed mRNAs and 651 significantly differentially expressed lncRNAs were detected. Furthermore, we identified 10 key genes that may affect reproductive performance in chickens: STC1, MMP13, IL8, AT2, FSHB, RARB, THOC7, FGF12, EPHA1, and NPY5R. Based on GO and KEGG enrichment analyses, we preliminarily explored five pathways that may influence egg production performance: neuroactive ligand-receptor interaction, CCR6 chemokine receptor binding, G-protein coupled receptor binding, cytokine-cytokine receptor interaction and ECM-receptor interaction. These genes and pathways may play crucial roles in improving broodiness behavior. This study establishes a foundation for subsequent screening of candidate functional genes and functional validation of important production traits in TBsf, providing a theoretical basis for further molecular breeding and enhancing egg-laying performance in TBsf.

## Introduction

Broodiness is a common habit of domestic fowls, which can inhibit egg production and affect the poultry industry [1]. Broodiness is observed in most breeds of domestic fowl, among which the black chicken shows a strong broodiness behavior. After every 15 eggs are laid, the broodiness occurs and the laying of eggs is stopped [2]. Broodiness is usually associated with increased body temperature, reduced feed and water intake, frequent nest occupancy, turning and retrieval of eggs, aggressive or defensive behaviors, characteristic clucking, and cessation of egg production [3]. Meanwhile, it is also associated with hereditary, nutrition, hormones, and so on. Genetic factors are the root cause of broodiness in fowls.

The broodiness behavior of poultry is highly regulated by the hypothalamic-pituitary-gonad (HPG) axis, and the changing tide of ovarian signals appears to determine to a large extent the nature of the activities of the hypothalamic-pituitary unit [4]. The ovary is a reproductive organ of vertebrates and the site of production and daily release of oocytes. The ovary consists of follicles at several different stages of development. Follicles are the basic unit of reproduction, which is composed of a central oocyte and the surrounding endocrine cells [5]. The domestic fowl is a unique model for studying follicular development. Unlike mammals, the single left ovary of the hen contains follicles of different sizes and development stages. The resting primordial follicles, pre- hierarchical growing follicles, pre-ovulatory follicles and post-ovulation follicles can appear simultaneously in an ovary with reproductive activity. Before final ovulation, the size of follicles can increase from 6-8 mm to 40 mm within 5-9 days [6]. Starting from the ovary will help us to explore the molecular mechanism that affects hen broodiness, and further provide theoretical basis for improving egg production.

The development of ovary and follicle is an important factor affecting the laying performance of chicken. The biological processes of ovarian development and ovulation are regulated by a large number of dynamic and stage-specific expression of key genes [7]. Therefore, studying the expression characteristics of chicken ovarian tissue during broodiness at the transcriptional level can provide an important basis for screening and identifying key genes that regulate chicken ovarian development. Given the rapid development of high-throughput sequencing, the analyses of transcriptomics have been used to screen out the candidate genes, pathways and molecular mechanism of broodiness.

Transcriptome sequencing is an effective method for investigating the important functions of long non-coding RNAs (lncRNAs), RNA transcripts >200 nucleotides that do not encode protein are broadly defined as lncRNAs [8,9]. LncRNAs may be involved in mRNA splicing and maturation, mRNA transport or localization, mRNAs stabilization [10].

We focus on these >200-nt-long RNA transcripts in ovaries with described noncoding functions. LncRNAs are recognized as playing crucial roles in reproductive processes, including sexual maturation [11], spermatogenesis [12], premature ovarian insufficiency [13]. Wang et al. identified 1182 mRNAs and 168 lncRNAs that differ in granulosa cells of small yellow follicles from Jinghai Yellow chickens in red light and white light groups, many involved in follicular development through steroid hormone synthesis, oocyte meiosis, and the PI3K-Akt signaling pathway [14]. Meanwhile, Wu et al. constructed lncRNA-gene interaction networks of 34 differen tially abundant lncRNAs and 263 DEGs, leading to an enhanced understanding of lncRNA and gene interactions regulating broodiness [15]. Taken together, the integrated analysis of lncRNA and mRNA may provide novel clues for exploring the mechanism that regulate broodiness.

The Taihe black-bone silky fowl (Gallus gallus domesticus Brissonis) native to Wangbantu village, Taihe county, Jiangxi province. It has the characteristics of “tassel head, tufted crown, green ear, beard, silk hair, black skin, black meat, black bone, hairy foot, and five claws”. It has high edible, medicinal and ornamental value, and is a valuable variety resource and traditional precious Chinese medicine in China[16,17]. The TBsf has been highly valued as a curative food with many healthcare function and pharmacological effects, such as anti-oxidative and anti-fatigue effects [16]. Broodiness is an instinct for breeding offspring, with TBsf having a strong broodiness behavior. Under natural conditions, TBsf generally broodiness once for every 10-12 eggs laid, each time for more than 15 days. This is extremely unfavorable for the economic development of TBsf industry. So far, Liao et al. have explored the muscle metabolites of TBsf influenced by the breed and feed traits at the metabolome level [18]. Xiang et al. have revealed the differentially expressed genes and pathways that affect egg production performance at the transcriptome level. However, the roles of lncRNAs in the reproduction of TBsf are unclear, and the impact of lncRNAs and mRNAs on ovarian development is still worth further research.

In the recent study, we selected two periods to explore the impact of broodiness on the laying rate of hens: the broodiness period and the peak egg-laying period. RNA sequencing was employed to compare ovarian lncRNAs and mRNAs in different periods. By comparing the two periods, we screened out the differentially abundant lncRNAs and mRNAs. LncRNAs were selected for bioinformatic analysis to predict cis- and trans-target genes to construct related networks of lncRNA genes. Subsequently, function and pathway enrichment analysis of important genes were conducted by Gene Ontology (GO) and Kyoto Encyclopedia of Genes and Genomes (KEGG). The results could prove useful for exploring the molecular mechanisms of broodiness in black chickens, and help to improve the egg laying performance of to them.

## Results

### RNA sequencing and mapping

There were 13.95GB of clean data from eight cDNA libraries (groups BC and GC), including 92,978,840 reads that were generated after quality control assessment with the filtering out of reads containing adaptor contamination, undetermined bases, and low-quality bases. The Q20 content ranged from 96.86% to 97.46%, the Q30 content ranged from 91.57% to 92.80%, and the GC content ranged from 46.61% to 47.62%. More than 94% of clean reads were mapped to the reference genome of chicken and then used for further gene expression analysis. Approximately 90% of the reads were uniquely aligned, and approximately 3.9% were aligned in multiple mapped pathways (Table 1).

**Table 1.** Summary statistics for sequence quality and alignment information.

| Sample | Raw reads | Clean reads | Raw bases (G) | Clean bases (G) | Q20 (%) | Q30 (%) | GC content (%) | Total mapped | Multiple mapped | Uniquely mapped |
| --- | --- | --- | --- | --- | --- | --- | --- | --- | --- | --- |
| BC1 | 94284862 | 92324972 | 14.14 | 13.85 | 97 | 91.9 | 47.07 | 86516681 (93.71%) | 3432068 (3.72%) | 83084613 (89.99%) |
| BC2 | 92798320 | 90386260 | 13.92 | 13.56 | 96.99 | 91.93 | 47.62 | 85059842 (94.11%) | 3983147 (4.41%) | 81076695 (89.7%) |
| BC3 | 106996926 | 104642326 | 16.05 | 15.7 | 97.46 | 92.8 | 47.25 | 99329692 (94.92%) | 4575356 (4.37%) | 94754336 (90.55%) |
| BC4 | 90841188 | 89148020 | 13.63 | 13.37 | 96.86 | 91.61 | 46.62 | 83569009 (93.74%) | 3511408 (3.94%) | 80057601 (89.8%) |
| GC1 | 92547334 | 90696320 | 13.88 | 13.6 | 97.06 | 92.09 | 47.45 | 85154842 (93.89%) | 3469965 (3.83%) | 81684877 (90.06%) |
| GC2 | 86443334 | 83522870 | 12.97 | 12.53 | 96.88 | 91.57 | 46.61 | 78938778 (94.51%) | 2221008 (2.66%) | 76717770 (91.85%) |
| GC3 | 103790790 | 101415404 | 15.57 | 15.21 | 97.12 | 92.21 | 47.15 | 95446885 (94.11%) | 3877094 (3.82%) | 91569791 (90.29%) |
| GC4 | 93705468 | 91694550 | 14.06 | 13.75 | 97.21 | 92.38 | 47.38 | 86472645 (94.31%) | 4233684 (4.62%) | 82238961 (89.69%) |

### Transcriptome assembly

In total, the assembled transcripts comprised 21,425 lncRNAs, including 8,870 known and 12,555 novel lncRNAs. The CPC2, CNCI, and Pfam tools were employed to assess the protein-coding potential of the transcripts in this study. Regarding the genomic locations of novel lncRNAs, 7,677 were intergenic (61.1%), 2,490 were antisense lncRNAs (19.8%) and 2,388 were sense-overlapping lncRNAs (19.0%) (Fig 1A).

**Fig 1.**
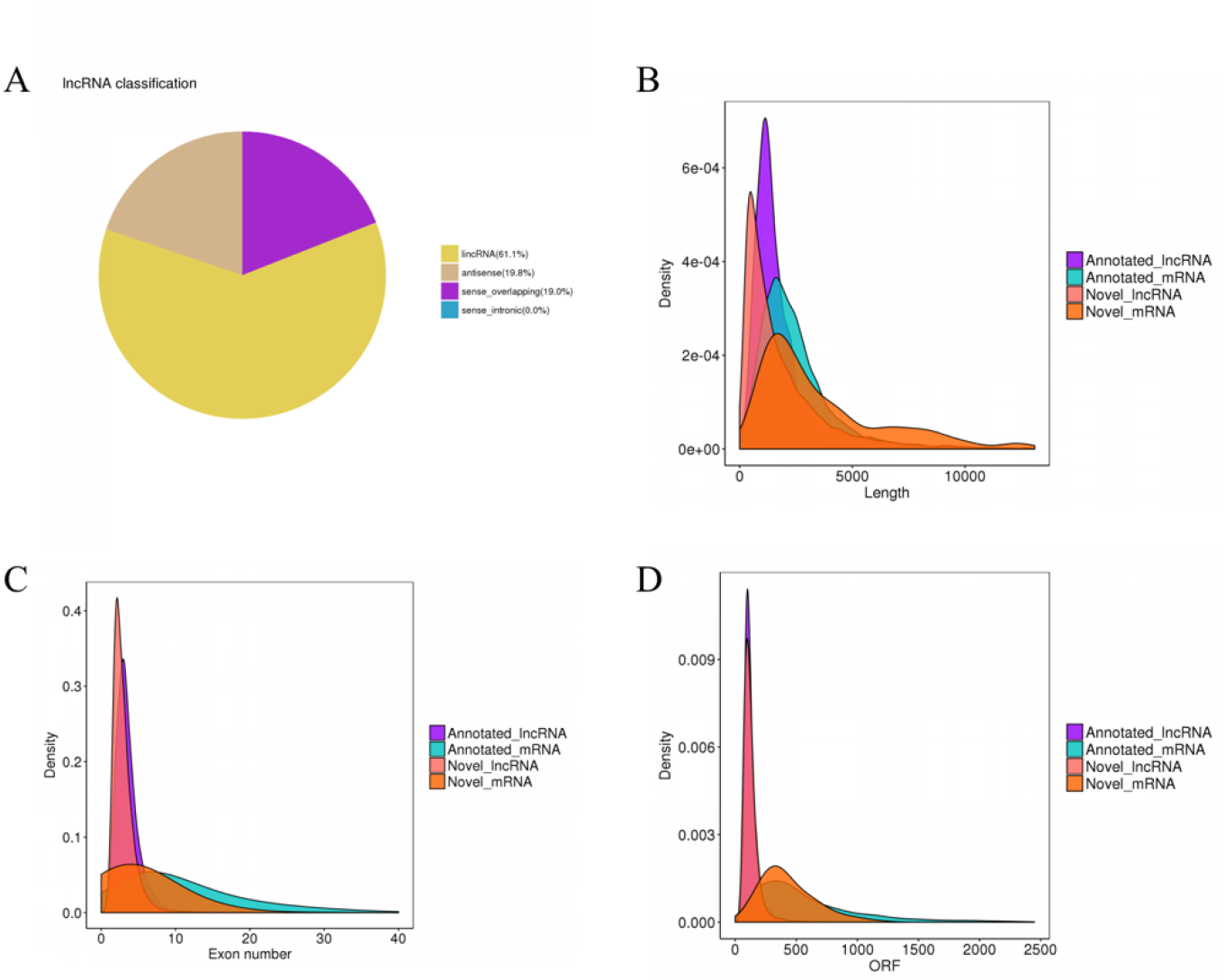
LncRNA classification and genomic features in the ovaries of TBsf. (A) LncRNA classification. (B) The transcript length distribution of lncRNAs and mRNAs. (C) The exon number distribution of lncRNAs and mRNAs. (D) The ORFs length distribution of lncRNAs and mRNAs.

In this study, the average length of lncRNA transcripts was 2049 bp, which is shorter than the 5607 bp length of mRNA transcripts, thus indicating that lncRNAs were shorter than mRNAs (Fig 1B). Additionally, the number of exons in lncRNAs (3.05 on average) was less than that of mRNAs (5.03 on average) (Fig 1C). Moreover, lncRNAs tended to have shorter ORFs than mRNAs in the ovaries (Fig 1D).

### Differentially expressed mRNAs and lncRNAs

The FPKM values of mRNAs and lncRNAs in the ovarian tissues from hens in the BC and GC groups were calculated (S1 and S2 Tables). After calculating the FPKM of all transcripts in each sample, the distribution of the expression level of different transcripts in the sample is displayed through the Boxplots (Fig 2A). To verify the repeatability between the groups and the correctness of the groups, we performed PCA. The PCA result showed that there was a certain distance between group BC and group GC, which indicates that the samples of group BC and group GC are different.

**Fig 2.**
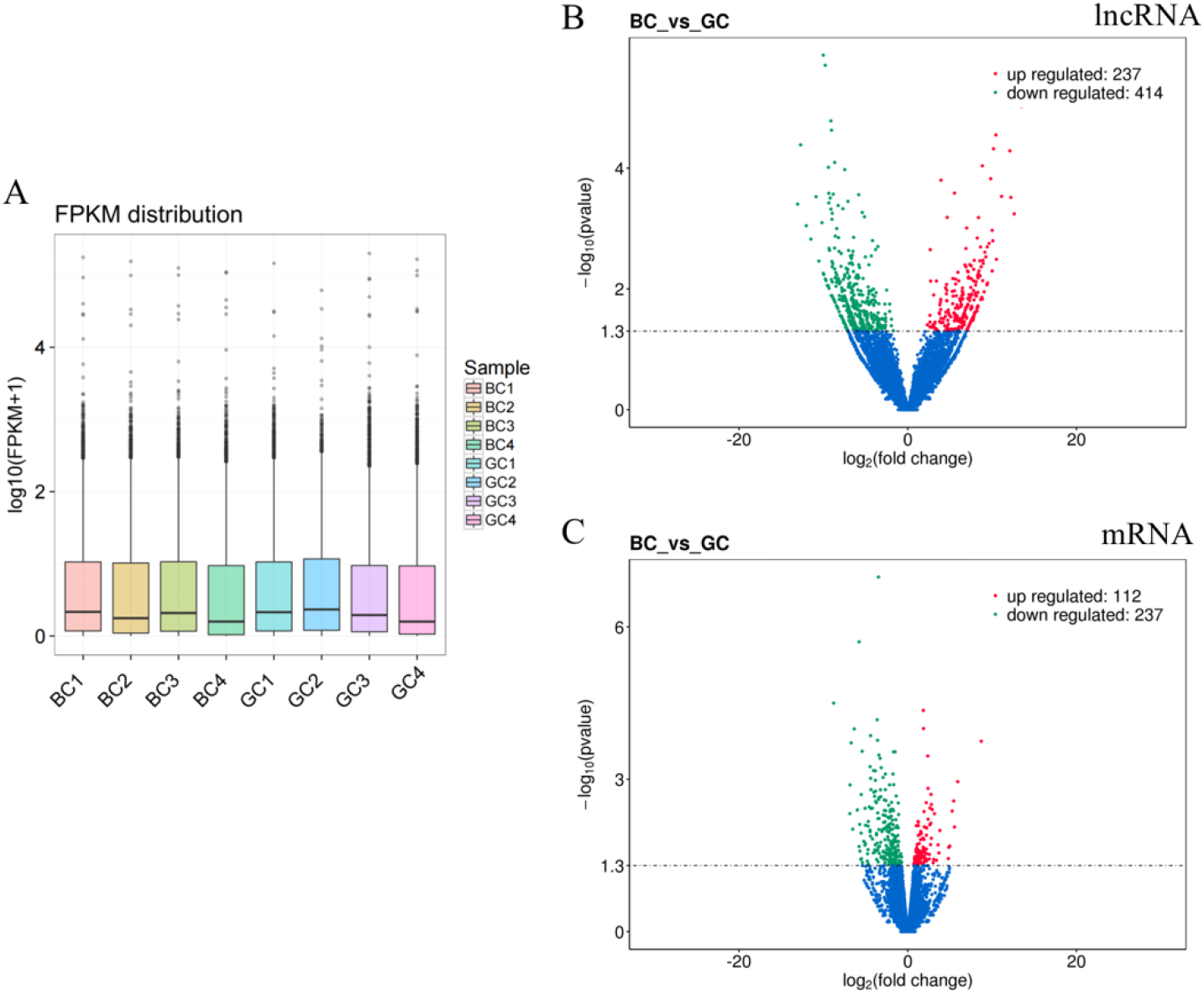
Comparison of expression level and the differential expression of lncRNAs and mRNAs between the BC and GC. Up-regulated genes are shown in red, down-regulated genes are shown in green, and genes with no significant difference in expression are indicated in blue; significance was indicated by a p-value < 0.05. (A) Boxplots of each sample expression levels. (B) Differential expression of lncRNAs. (C) Differential expression of mRNAs.

To identify the DE lncRNAs and mRNAs between the BC and GC groups, we used edgeR software. As a result, 349 mRNAs and 651 lncRNAs were found to be significantly differentially expressed between the BC and GC groups fowl ovaries. 237 lncRNAs and 112 mRNAs were significantly upregulated, while 414 lncRNAs and 237 mRNAs were downregulated (Fig 2B, C).

### Enrichment analysis of differentially expressed mRNAs

GO is a comprehensive database describing gene functions, which can be divided into three parts: biological processes (BPs), cellular components (CCs), and molecular functions (MFs). In the BC vs. GC groups, a total of 321 differentially expressed mRNAs were enriched to 4850 GO terms with functional annotation information, including 3745 BPs, 392 CCs, and 713 MFs. There were 797 GO terms significantly enriched in the GO results that met the criteria of P < 0.05 (S3 Table). The significantly enriched terms included extracellular region, chemokine receptor binding, multi-organism process, CCR6 chemokine receptor binding and G-protein coupled receptor binding (Fig 3A). KEGG pathway analysis recognized 16 significantly enriched pathways (P < 0.05), including ECM-receptor interaction, neuroactive ligand-receptor interaction, Cytokine-cytokine receptor interaction, NOD-like receptor signaling pathway and Focal adhesion (Fig 3B and S4 Table).

**Fig 3.**
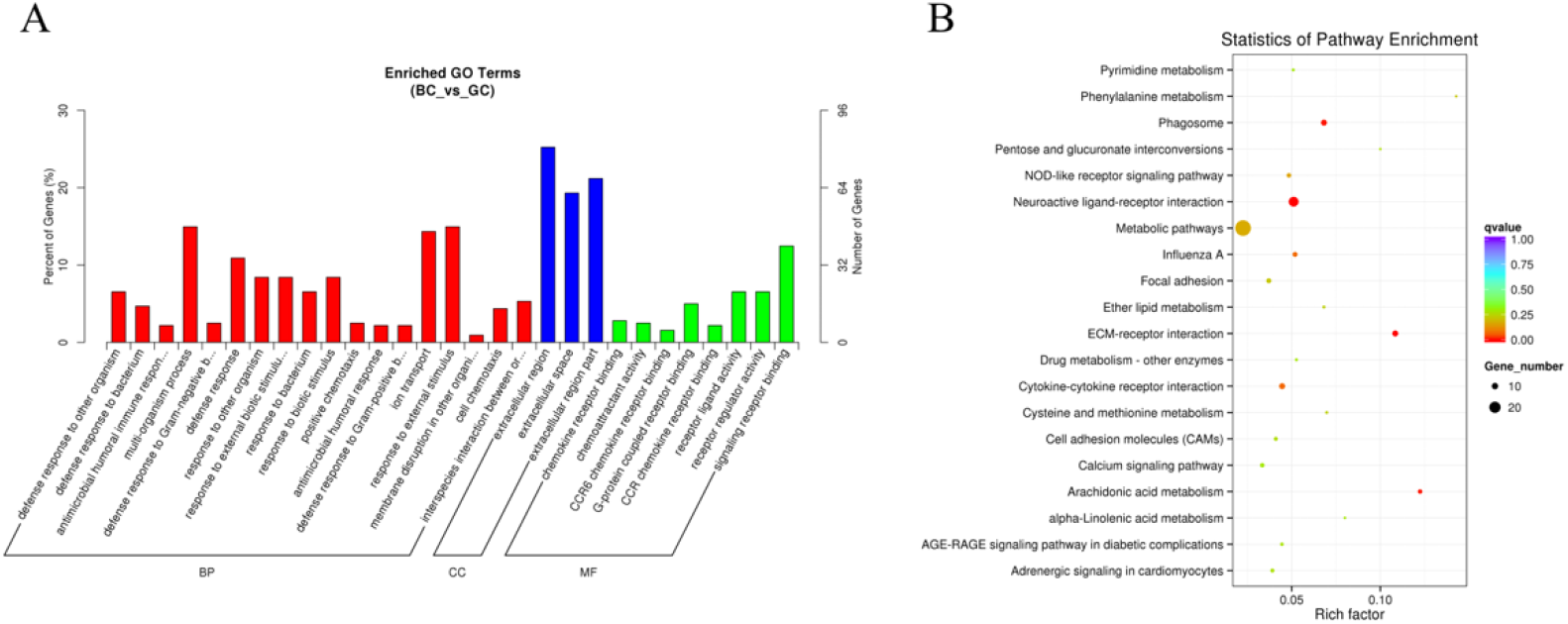
GO and KEGG analysis of differential mRNA expression. (A) Histogram of GO enrichment of DE mRNAs. (B) Scatter plot of KEGG enrichment for DE mRNAs.

### Regulatory roles of differentially expressed lncRNAs in ovaries

To explore the regulatory functions of lncRNAs, we predicted potential cis-acting and trans-acting targets of DE lncRNAs between the BC and GC groups. For cis-acting regulation of lncRNAs, we searched for protein-coding genes within 100 kb upstream and downstream of the lncRNAs. As a result, we identified 2,163 potential cis-regulated target genes (S5 Table). Based on these cis-regulated target genes, GO analysis revealed 804 significantly enriched GO terms (p < 0.05). The DE lncRNA target genes were found to be involved in activities related to CCR6 chemokine receptor binding, neuropeptide receptor activity, G-protein coupled peptide receptor activity, response to hormone and cell-cell signaling (Fig 4A and S6 Table). Pathway analysis indicated that these cis-regulated target genes of lncRNAs were enriched in 43 KEGG pathways, some of which were associated with ovarian follicle development, such as the Neuroactive ligand-receptor interaction, Progesterone-mediated oocyte maturation, Ribosome biogenesis in eukaryotes, Oocyte meiosis and MAPK signaling pathway (Fig 4B and S7 Table). These findings suggest that lncRNAs act in cis on neighboring protein-coding genes to regulate ovarian follicle development.

**Fig 4.**
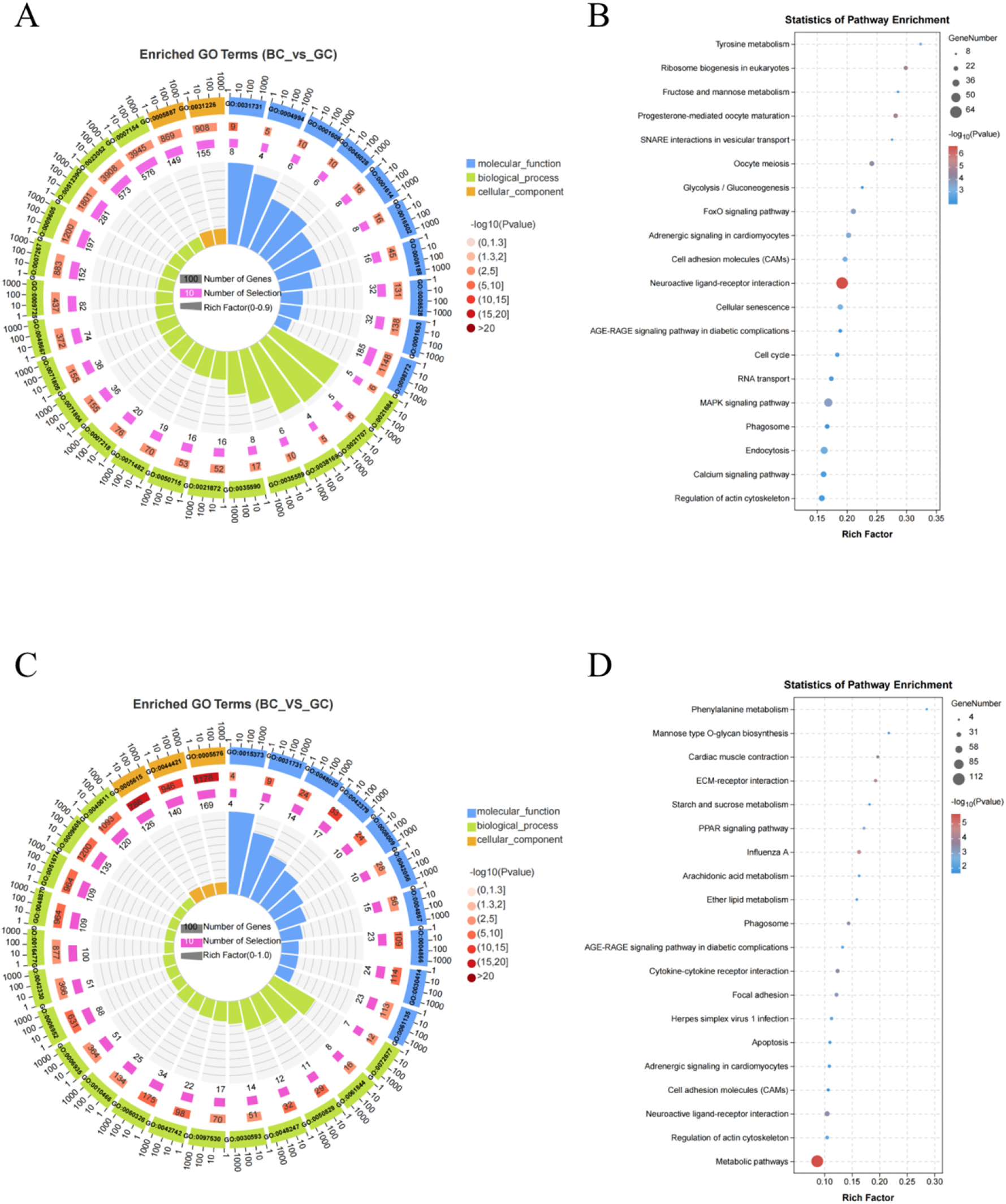
GO and KEGG analysis of differential lncRNAs target gene. (A) Circos plot of GO enrichment of target gene of DE lncRNAs in cis-regulatory. (B) Scatter plot of KEGG enrichment of target gene of DE lncRNAs in cis-regulatory. (C) Circos plot of GO enrichment of target gene of DE lncRNAs in trans-regulatory. (D) Scatter plot of KEGG enrichment of target gene of DE lncRNAs in trans-regulatory.

In the result of trans-regulated target genes of DE lncRNAs, we found 1302 potential lncRNA target genes (S8 Table). The lncRNAs from co-expression genes were significantly enriched in 638 GO terms (469 biological (BP), 47 cellular components (CC), and 122 molecular functions (MF)) that encompassed various biological processes. The main terms include extracellular region, chemokine receptor binding, defense response, response to external stimulus and cell motility (Fig 4C and S9 Table).

In addition, KEGG pathway analysis of differentially abundant trans lncRNAs was performed to identify pathways that were enriched by expression of these genes. The differentially abundant lncRNAs that were co-expression with protein-coding genes were enriched in 33 KEGG pathways, including metabolic pathways, ECM-receptor interaction, cytokine-cytokine receptor interaction, neuroactive ligand-receptor interaction and focal adhesion (Fig 4D and S10 Table). These findings suggest that lncRNAs act on protein-coding genes associated with ovarian follicle development in a trans-regulatory manner.

### Target gene prediction of lncRNAs and interaction network construction

To understand the biological mechanisms of putative lncRNAs in ovarian follicle development, we employed bioinformatics tools to construct a regulatory network for the putative lncRNAs and their potential targets. Since most lncRNAs regulate target genes in a trans-regulatory manner, we assembled a lncRNA-mRNA trans-regulatory interaction network associated with ovarian follicle development (Fig 5). Fig 5 illustrates the diverse regulatory relationships between the predicted lncRNAs and their trans-regulated target genes. Within the integrated network, 9 central node genes and 6 central node lncRNAs were identified. We found that TCONS_00017892 regulates a significant portion of the mRNAs within the network.

**Fig 5.**
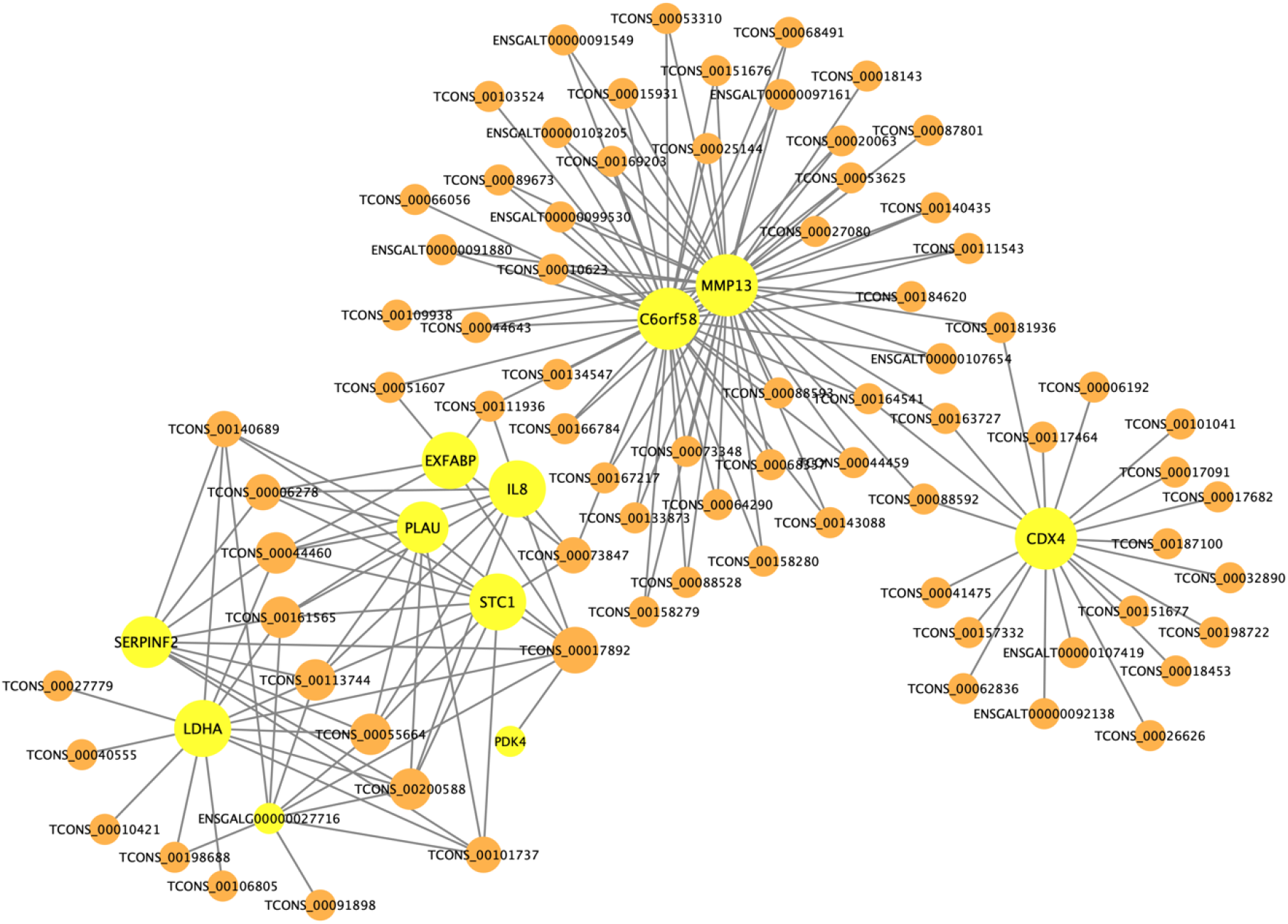
Interacting network of lncRNAs and their trans-regulated target genes associated with ovarian follicle development. The orange circles and yellow circles represent lncRNAs and potential target genes, respectively, with varying sizes based on their relative importance.

## Discussion

Egg production is an important index to evaluate the reproductive performance of hens. But hen’s nesting behavior can lead to the reduction of egg production. Therefore, it is of potential economic value to explore the behavior mechanism of chicken nesting for improving its egg production performance in genetics and breeding. Nesting is regulated by many critical genes and complex regulatory pathways. However, the mechanism underlying broodiness in TBsf is unclear. Egg production traits are related to ovarian function and are regulated by the hypothalamic-pituitary-gonadal (HPG) axis [19]. Therefore, ovarian tissue was selected to perform the RNA-seq analysis. At present, many studies have found that lncRNAs and mRNAs play critical roles in the regulation of ovary development. Ovary lncRNAs and mRNAs have been identified in chickens [11], Yili geese [20], Hu sheep [21], Muscovy ducks [15], domestic pigeons [22] and Chinese Tongue Sole [23]. Although the mRNA transcriptome in TBsf ovary has been partially explored in previous studies [24], a systematic evaluation of the lncRNAs and mRNAs across TBsf ovary developmental stages and broodiness stages is lacking. So, in this study, we constructed 8 cDNA libraries from the ovaries of BC and GC groups to evaluate the expression of mRNAs and lncRNAs by Illumina high-throughput sequencing, and we identified critical candidate lncRNAs related to broodiness. Specifically, we identified 349 DE mRNAs and 651 DE lncRNAs between BC and GC groups fowl ovaries. In addition, we found that lncRNAs identified in this study have shorter transcript lengths, fewer exons and shorter ORFs than protein coding transcripts, which indicated that the lncRNAs were of high quality and that the data collected were reliable.

In this study, we integrated the interaction network of differentially expressed lncRNAs and their trans-regulated target genes to identify several potential mRNAs associated with chicken ovarian development, including STC1, MMP13 and IL8. Stanniocalcin-1 (STC-1) is a widely expressed glycoprotein hormone involved in various physiological and pathological processes, including angiogenesis, mineral homeostasis, cell proliferation, inflammation, and apoptosis. Within female reproductive tissues, STC-1 expression is associated with processes such as ovarian follicle development, embryonic implantation, early pregnancy vascular remodeling, and placental development [25]. STC-1 is recognized as a crucial ovarian regulatory factor, promoting ovarian development in mice. It stimulates granulosa cell proliferation and affects the secretion of E2 (estradiol) and P4 (progesterone) by influencing the expression of key genes involved in steroid hormone synthesis [26]. Metalloproteinases (MMPs) are present in the chicken ovary and play functional roles in regulating the extracellular matrix (ECM) during processes such as follicular development, ovulation, atresia, and regression [27]. MMP13, also known as collagenase-3, initiates the degradation of fibrillar collagen, a critical component in the formation of key structural elements of membranes. In chickens, MMP13 is involved in embryonic membrane remodeling and is associated with biochemical and molecular changes related to corneal development. In laying hens and POF1 (newly postovulatory follicles) ovarian follicles, both the mRNA and protein expression of chicken MMP13 are significantly upregulated, with the chicken MMP13 protein primarily expressed in sexually mature ovarian membrane cells [28]. Interleukin-8 (IL-8), also known as CXCL8, is a well- known chemotactic cytokine secreted by various cells. Recent studies have demonstrated the involvement of IL-8 in the establishment of the corpus luteum following ovulation. IL-8 contribute to angiogenesis and IL-8 specifically stimulates progesterone secretion through its actions on luteinized granulosa cells and luteal membrane cells [29]. IL-8 also participates in the regulation of inflammatory responses and immune modulation in ovarian tissues. Aberrant expression of IL-8 may be associated with the occurrence and progression of ovarian diseases, including polycystic ovary syndrome and ovarian tumors [30,31].

Furthermore, we also identified genes within the differentially expressed gene set that may have potential associations with broodiness in hens: AT2 and FSHB. Angiotensin II (Ang II) is a key component of the renin-angiotensin system (RAS) and is known to have diverse physiological functions, including blood pressure regulation, fluid homeostasis, and vascular smooth muscle proliferation. The involvement of RAS in the synthesis and secretion of prohormones and estrogens, as well as the regulation of ovarian follicle development, ovulation, and luteal regression. The AT2 receptor is believed to play a role in follicular cell communication during ovarian luteal regression [32]. The presence of Angiotensin II (Ang II) in porcine follicular fluid (pFF) and porcine ovaries suggests its significant involvement in the regulation of oocyte maturation and follicular development. Ang II likely plays a crucial role in modulating ovarian function by regulating intrafollicular estrogen concentrations, which are vital for maintaining dominant follicles [33]. FSHB, the beta subunit of follicle-stimulating hormone (FSH), combines with the alpha subunit (FSHA) to form biologically active FSH. FSH plays a crucial role in reproductive competence. By binding to receptors on ovarian cells, FSH regulates the development and maturation of follicles. It stimulates granulosa cell proliferation, facilitates estrogen synthesis and secretion, and promotes the development of oocytes within the follicles. FSH is also involved in the regulation of corpus luteum formation and progesterone synthesis [34,35]. Among them, IL8 is enriched in the signaling pathways of cytokine-cytokine receptor interaction and G-protein coupled receptor binding. On the other hand, FSHB is enriched in the signaling pathway of neuroactive ligand-receptor interaction.

Through analysis of differentially expressed lncRNAs, we identified several potential cis-regulated target genes that may be involved in ovarian function regulation, including RARB, THOC7, FGF12, EPHA1, and NPY5R. Interestingly, THOC7 also emerged as a trans-regulated target gene. Retinoic acid (RA) is an effective inducer of cellular differentiation and plays a crucial role in the development of sexually dimorphic reproductive cells in mammals. RA is known to be involved in cellular differentiation and maintenance of cell fate in somatic cells of the ovary. Retinoic Acid Receptor Beta (RARB) is a member of the retinoic acid receptor family. As a nuclear receptor, RARB participates in the regulation of gene transcription, primarily through its involvement in the retinoic acid signaling pathway [36]. RARB exhibits significant expression in the ovary, particularly in granulosa cells and oocytes. However, there is limited or no research available on the involvement of the three RXR protein isoforms (RXRα, RXRβ, RXRγ) in ovarian function. Further investigations are necessary to elucidate the precise role and molecular mechanisms of RARB in the ovary [37]. The THO complex, a multi-subunit family with transcriptional and mRNA export functions, plays an indispensable role in embryogenesis, organogenesis, and cell differentiation. Differential gene expression of THOC in the ovary suggests its specific involvement in follicle development and oogenesis processes. The higher expression of THOC7 within these genes implies its potential significant role in ovarian steroidogenesis or regulation of other factors that govern ovarian growth. THOC7 may therefore be implicated in ovarian growth and maturation [38]. FGF12, a growth factor, exerts various effects including promoting endothelial cell migration, smooth muscle cell proliferation, new blood vessel formation, and facilitating repair of damaged endothelial cells. It acts intracellularly by inhibiting cellular apoptosis. Downregulation of FGF12 may also potentially contribute to the apoptosis of closed follicles. Within female reproductive tissues, FGF12 exhibits high expression specifically in the ovary. Its expression facilitates ovarian development while suppressing premature cellular apoptosis. Research suggests that FGF12 expression promotes the absorption of fatty acids in the yolk sac, leading to increased body weight and enhanced ovarian development. Therefore, FGF12 emerges as a potential candidate gene involved in regulating follicular closure [39,40]. The Eph family is the largest subgroup within the receptor tyrosine kinase family, consisting of EphA (EphA1-10) or EphB (EphB1-6) receptor subclasses. Eph receptor-A1 (EphA1) is the first member of the Eph receptor tyrosine kinase family implicated in erythropoietin-producing hepatocellular carcinoma (Eph). Many members of the Eph family and their ligands have been identified in ovarian tissue, participating in ovarian development, growth, and functional regulation. These studies suggest a potential role for the Eph family in the ovaries. For instance, ephrin- B1 and EphB4 are co-expressed in all types of steroid-producing cells present in the ovary [41]. Furthermore, aberrant expression of EphA2 receptor and EphrinA1 ligand is associated with the occurrence and progression of ovarian cancer. EphA1 may function as a tumor suppressor in ovarian cancer [42,43]. Although direct evidence linking the EPHA1 gene to the ovaries is limited, understanding the significance of the Eph family in ovarian biology provides a deeper insight into ovarian-related diseases and treatments. Neuropeptide Y receptor type 5 (NPY5R) is a G-protein coupled receptor belonging to the neuropeptide Y (NPY) receptor subfamily, mediating the effects of endogenous NPY. Research has shown that NPY5R is involved in regulating the proliferation and apoptosis of granulosa cells, thereby modulating female reproductive functions through the central nervous system. NPY5R signaling is likely to be an intraovarian regulatory event in folliculogenesis, promoting the survival and growth of early antral follicles [44,45]. Among these genes, NPY5R is enriched in the pathways of neuropeptide receptor activity and G-protein coupled peptide receptor activity.

We found that numerous lncRNAs target genes and mRNAs were both involved in the regulation of neuroactive ligand-receptor interaction, CCR6 chemokine receptor binding, G-protein coupled receptor binding, Cytokine-cytokine receptor interaction and ECM-receptor interaction. Neuroactive ligands affect neuronal function by binding to intracellular receptors, which have the capability of binding transcription factors and regulating gene expressions [46]. Mu et al.’s study suggested neuroactive ligand-receptor interaction pathway might affect egg production in chickens via a mechanism similar to that found in fish [47]. Caballero-Campo et al. demonstrated that CCR6 protein is localized on the surface of human sperm [48]. Chemokines play various biological functions by activating surface receptors of their target cells, and also have the ability to interfere with sperm–oocyte interaction [49]. In addition, CCR6 specifically binds to its ligand chemokine CCL20. Duan et al. demonstrated that the chemokine CCL20 is abundantly present in human follicular fluid and is produced by human oocytes as well as surrounding cumulus granulosa cells [50]. This chemokine has an important function for the process of fertilization. G protein-coupled receptors (GPCRs), representing the largest protein family encoded by the human genome, are membrane-bound receptors that mediate crucial physiological responses by converting extracellular signals. These receptors exhibit a diverse array of endogenous ligands, encompassing odorants, hormones, neurotransmitters, and chemotactic factors. The ligands for GPCRs span a broad spectrum, ranging from photons, amines, carbohydrates, lipids, peptides, to proteins [51]. In the ovary, multiple GPCRs play a role in regulating reproductive functions. These receptors interact with endogenous hormones, neurotransmitters, or other signaling molecules within the ovary, thereby influencing important physiological processes such as ovarian development, follicle growth, and ovulation. Investigation of these GPCR signaling pathways contributes to a better understanding of the regulatory mechanisms underlying ovarian function. Cytokines are soluble extracellular proteins or glycoproteins that play critical roles as intercellular mediators and mobilizers in innate and adaptive immune host defense, cell growth, differentiation, cell death, angiogenesis, and developmental and reparative processes aimed at restoring homeostasis. They are released by various cells in response to activating stimuli and induce specific biological responses by binding to specific receptors on the cell surface of target cells. Existing research has provided evidence supporting the significant involvement of the cytokine-cytokine receptor interaction pathway in follicle development [52]. The extracellular matrix (ECM) is a complex matrix of biomacromolecules, including glycoproteins, proteoglycans, and glycosaminoglycans. The ECM-receptor interaction pathway is the most significantly enriched signaling pathway in terms of gene enrichment. It plays a pivotal role in various aspects of cellular physiological activities, such as cell adhesion, migration, proliferation, and differentiation [53]. Transcripts associated with the “ECM-receptor interaction” pathway may complement the enrichment of Gene Ontology categories related to cell adhesion in biological processes, indicating their potential significance in promoting cell adhesion and cohesion. This evidence underscores the crucial molecular-level importance of adhesion and cohesion mechanisms in the physiological activities of the ovary [54].

## Materials and methods

### Ethical statement

All animal experiments conformed to the standards in the Chinese animal welfare guidelines and were approved by the Animal Experimentation Ethics Committee of Zhejiang University (approval number: ZJU20190149).

### Animal and sample collection

Eight 30-week-old female TBsf (comprising four broodiness chickens (BC) and four high egg-laying chickens (GC)) were purchased from the Taihe county in the Jiangxi province from the Taihe Aoxin black-bone silky fowl Development Co (Taihe county, Jiangxi province, China).

To classify TBsf on the basis of brooding and high egg-laying periods, eggs were continuously collected to record the egg-laying patterns. The chickens were fed according to the same housing and feeding conditions. All chickens were humanely slaughtered and necropsied. The tissues of the entire ovary were collected, frozen in liquid nitrogen, and stored at −80 ◦C until further manipulation.

### RNA isolation, library construction and sequencing

The collected hen ovarian tissues were delivered to the Novogene Bioinformatics Technology Co., Ltd. (Beijing, China), who conducted all the library preparation and sequencing. Total RNA was isolated from each sample. The integrity of RNA and whether there was DNA contamination was assessed using agarose gel electrophoresis. The purity and concentration of total RNA were determined using NanoDrop procedures. Using Agilent 2100 bioanalyzer can accurately detect RNA integrity.

Ribosomal RNA (rRNA) was removed from the total RNA, and then the RNA was broken into short fragments of 250 to 300 bp. The first strand of cDNA was synthesized using fragmented RNA as a template and random oligonucleotides as primers, followed by the synthesis of a second strand of cDNA using dNTPs (dUTP, dATP, dGTP, and dCTP) as raw materials. The purified double stranded cDNA was subjected to terminal repair, A-tailed addition, and sequencing adapter were connected. In order to select cDNA fragments of preferentially 350∼400 bp in length, the library fragments were purified with AMPure XP bead. Using the USER enzyme (NEB, California, USA) to degrade the second strand of U-containing cDNA, PCR amplification was performed and libraries was obtained. The library quality was assessed using Agilent Bioanalyzer 2100 system and the libraries were sequenced on an Illumina NovaSeq 6000.

### Bioinformatics analysis

First, raw reads in FASTQ format were processed with an in-house Perl script. In order to ensure the quality and reliability of data analysis, it is necessary to obtain clean reads by removing low-quality reads containing adaptor, poly-N sequences and low-quality bases from the raw data and calculating the Phred score (Q20 and Q30) and GC content. All subsequent analyses were based on readings with reliable data. Paired-end clean reads were mapped with Hisat2 [55] to the reference genomes (http://ftp.ensembl.org/pub/release-105/fasta/gallus_gallus/, http://ftp.ensembl.org/pub/release-105/gtf/gallus_gallus/) to align all reads to obtain as small a transcript collection as possible. StringTie was used to splice and quantify the reads into transcripts based on the results of the comparison to the genome [56]. Cuffmerge was used to merge the transcripts obtained from splicing various samples and to remove transcripts with uncertain chain direction and transcript length not exceeding 200 nt. Fragments per kilobase of transcript per million mapped reads (FPKM) values were used to calculate the expression levels of the transcripts using the StringTie. After quantitative analysis, edgeR was used to identify differentially expressed lncRNAs and mRNAs. The differentially expressed mRNAs and lncRNAs were identified as having a statistical significance (p value < 0.05).

### Co-expression (trans) and co-location (cis) analyses

Cis-acting lncRNAs target neighboring protein-coding genes, and the protein-coding genes located within 100-kb upstream and downstream of the lncRNA are classified as cis-regulated target genes. To classify trans-regulated target genes of lncRNAs, we analyzed the abundance of identified lncRNAs and known protein-coding genes in laying and brooding chickens to understand the correlation between lncRNA abundance and protein-coding gene expression. We calculated the Pearson correlation coefficients (|r| > 0.95) between lncRNA and mRNA for further analysis. Additionally, we used Cytoscape v3.10.0 software to select differentially abundant lncRNAs and their corresponding differentially expressed cis-regulated and trans-regulated target genes to construct an lncRNA-gene interaction network.

### GO and KEGG enrichment analysis

To obtain further insight into the functions and classifications of differentially expressed genes and lncRNA target genes, we performed Gene Ontology (GO) term and Kyoto Encyclopedia of Genes and Genomes (KEGG) pathway analyses using GOseq and KOBAS software. GO is a comprehensive database describing gene functions, which can be divided into three parts: molecular function, biological process, and cellular component. KEGG is a comprehensive database integrating genomic, chemical, and system functional information [57]. A P-value<0.05 was considered to be a significant enrichment of differentially expressed genes.

## Supporting information

**S1 Table. The FPKM values of mRNAs.**

(XLS)

**S2 Table. The FPKM values of lncRNAs.**

(XLS)

**S3 Table. GO enrichment result of DE mRNAs.**

(XLS)

**S4 Table. KEGG pathway enrichment result of DE mRNAs.**

(XLS)

**S5 Table. Cis-regulated target genes of DE lncRNAs.**

(XLS)

**S6 Table. GO enrichment analysis of cis-regulated target genes of DE lncRNAs.**

(XLS)

**S7 Table. KEGG pathway enrichment analysis of cis-regulated target genes of DE lncRNAs.**

(XLS)

**S8 Table. Trans-regulated target genes of DE lncRNAs.**

(XLS)

**S9 Table. GO enrichment analysis of trans-regulated target genes of DE lncRNAs.**

(XLS)

**S10 Table. KEGG pathway enrichment analysis of trans-regulated target genes of DE lncRNAs.**

(XLS)

## Author Contributions

**Conceptualization:** Yuting Tan, Yunyan Huang, Zhaozheng Yin.

**Data curation:** Yuting Tan, Yunyan Huang.

**Formal analysis:** Yuting Tan, Yunyan Huang, Chunhui Xu, Xuan Huang.

**Funding acquisition:** Zhaozheng Yin.

**Investigation:** Chunhui Xu, Xuan Huang, Zhaozheng Yin.

**Resources:** Yunyan Huang, Chunhui Xu.

**Software:** Yuting Tan, Xuan Huang.

**Visualization:** Yuting Tan, Yunyan Huang.

**Writing –original draft:** Yuting Tan, Yunyan Huang, Chunhui Xu, Xuan Huang, Zhaozheng Yin.

**Writing –review & editing:** Yuting Tan, Yunyan Huang, Chunhui Xu, Xuan Huang, Zhaozheng Yin.

## Notes

### Competing Interest Statement

The authors have declared no competing interest.

